# HES1 contributes to high salt stress response as an enhancer of NFAT5-DNA binding

**DOI:** 10.1101/2022.10.10.511678

**Authors:** Hiroki Ryuno, Yusuke Hanafusa, Isao Naguro, Hidenori Ichijo

## Abstract

High salt conditions and subsequent hyperosmolarity are injurious cellular stresses but can activate immune signaling. Nuclear factor of activated T-cells 5 (NFAT5) is an essential transcription factor that induces osmoprotective genes such as aldose reductase (AR) and betaine-GABA transporter 1 (BGT1). High salt stress-mediated NFAT5 activation is also reported to accelerate the inflammatory response and autoimmune diseases. However, the systemic regulation of NFAT5 remains unclear. Here, we performed a genome-wide siRNA screen to comprehensively identify the upstream factors of NFAT5. We monitored NFAT5 nuclear translocation and identified one of the Notch signaling effectors, Hairy and enhancer of split-1 (HES1), as a novel positive regulator of NFAT5. HES1 was induced by high salinity via ERK signaling and facilitated NFAT5 recruitment to its target promoter region, resulting in the proper induction of osmoprotective genes and cytoprotection under high salt stress. These findings suggest that although HES1 is well known as a transcriptional repressor, it positively regulates NFAT5-dependent transcription in the context of a high salinity/hyperosmotic response.

**One Sentence Summary:** HES1 contributes to high salinity/hyperosmotic response through positive regulation of NFAT5-dependent transcription.

## INTRODUCTION

High salinity and consequential hyperosmotic conditions are environmental stresses commonly observed in our bodies. The stress physically induces water efflux from cells and subsequent cell shrinkage. Changes in intracellular molecule concentration and ion strength due to volume shrinkage cause DNA damage and protein misfolding (*1*). Cells manage such injurious stress by ion uptake to attenuate the difference between intercellular and extracellular osmolarity, resulting in short-term recovery of cellular volume. Then, cells absorb and synthesize osmolytes that are organic compounds, including sugars and amino acids, to replace ions to reduce intracellular ion strength after cell volume recovery (*2*). The accumulation of osmolytes is controlled by several enzymes and membrane transporters, such as aldose reductase (AR) and sodium- and chloride-coupled betaine transporter 1 (BGT1). Their expression are comprehensively regulated by a pivotal transcription factor, nuclear factor of activated T cells 5 (NFAT5; also called TonEBP/OREBP) (*3*).

NFAT5 is an essential transcription factor that responds to hyperosmotic conditions. NFAT5 is activated and translocates to the nucleus following stress and promotes the induction of osmoprotective genes such as regulators of osmolytes and protein chaperones, contributing to cellular homeostasis against hyperosmotic stress (*4*). Additionally, recent studies have reported that NFAT5 activation under high salt conditions skews immune cell differentiation toward a proinflammatory phenotype (*5, 6*). Although the phenomenon has beneficial aspects, such as antimicrobial defense and tumor suppression (*7, 8*), it also induces excessive inflammation under high salt intake, leading to the onset and progression of autoimmune and allergic diseases (*9, 10*). Thus, elucidation of the molecular mechanisms of NFAT5 activation caused by high salt stress is a promising way to identify a new therapeutic strategy for immune-related diseases.

To date, several upstream activators of NFAT5, such as ATM, CDK5 and cAbl1, have been individually reported under hyperosmotic conditions (*11*–*13*). However, a systematic understanding of NFAT5 regulation and coordination between pathways remains elusive. Here, we performed a genome-wide siRNA screen focusing on NFAT5 nuclear translocation under high salt conditions to comprehensively explore upstream regulators of NFAT5 and obtained 1291 candidate genes. A pathway analysis of the candidates revealed that genes in the Notch signaling pathway were enriched and that HES1, an effector of Notch signaling, strongly contributed to NFAT5 nuclear translocation and subsequent target gene induction. Although HES1 is known to be a transcriptional repressor, we revealed that HES1 enhances NFAT5 activity by facilitating the translocation of NFAT5 to the promoter region of a target gene. Furthermore, we found that HES1 is induced by high salinity and exhibits a cytoprotective effect under high salt stress. Our findings indicate that HES1 contributes to a high salinity/hyperosmotic response as a novel positive regulator of NFAT5.

## RESULTS

### HES1 is a novel regulator of NFAT5 nuclear translocation

We conducted a genome-wide siRNA screen to comprehensively elucidate the molecular mechanism of NFAT5 activation. In this screen, we monitored NFAT5 nuclear translocation, which is required for gene induction via NFAT5, under high salt conditions. We used HeLa cells stably expressing GFP in this screen for the high adherence under hyperosmotic conditions. The GFP signal was used for the definition of the whole-cell region (Fig. S1A). We introduced and stably expressed NFAT5 Δ547-tdTomato (NFAT5Δ), a deletion mutant that translocates to the nucleus by hyperosmolarity in the same way as full-length NFAT5 (*14*). In this study, the nuclear translocation ratio (TL ratio) of NFAT5 indicates the ratio of abundance in the nuclear region defined by the Hoechst signal to that in the whole-cell region defined by the GFP signal (Fig. S1A). The TL ratio of NFAT5Δ measured by a high-content image analyzer showed a linear increase in HeLa cells until three hours after high salt stimuli (Fig. S1B). Thus, we treated HeLa cells with additional 100 mM NaCl (500 mOsm) for three hours and evaluated the effect of each gene depletion on the TL ratio using a genome-wide siRNA library (>18000 genes) in the arrayed format.

In our screen, the known regulators of NFAT5 translocation, such as c-Abl1 and CDK5 (*11, 12*), had a relatively high rank (Sup Table 1), suggesting that our screen was successful in identifying regulators of NFAT5 translocation. We obtained 1291 genes whose knockdown attenuated NFAT5 nuclear translocation more strongly than the knockdown of c-Abl1 (TL ratio < 0.58) and defined these genes as the positive hits of the screen (Fig. 1A). A pathway analysis to explore key biological processes indicated significant enrichment of Notch signaling pathway-related genes in the positive hits (Fig. 1, B and C). Although the Notch signaling pathway is known to be important for developmental processes and cell differentiation (*15, 16*), little is known about the involvement of Notch signaling in the high salinity/hyperosmotic stress response. The positive hits included in the Notch signaling existed at various steps of the signaling pathway: a Notch receptor (Notch1), ligands (JAG1/2), gamma-secretase components (PSEN1/2 and PSENEN) and a downstream effector (HES1). Therefore, we focused on Notch signaling-related genes.

**Fig. 1.**
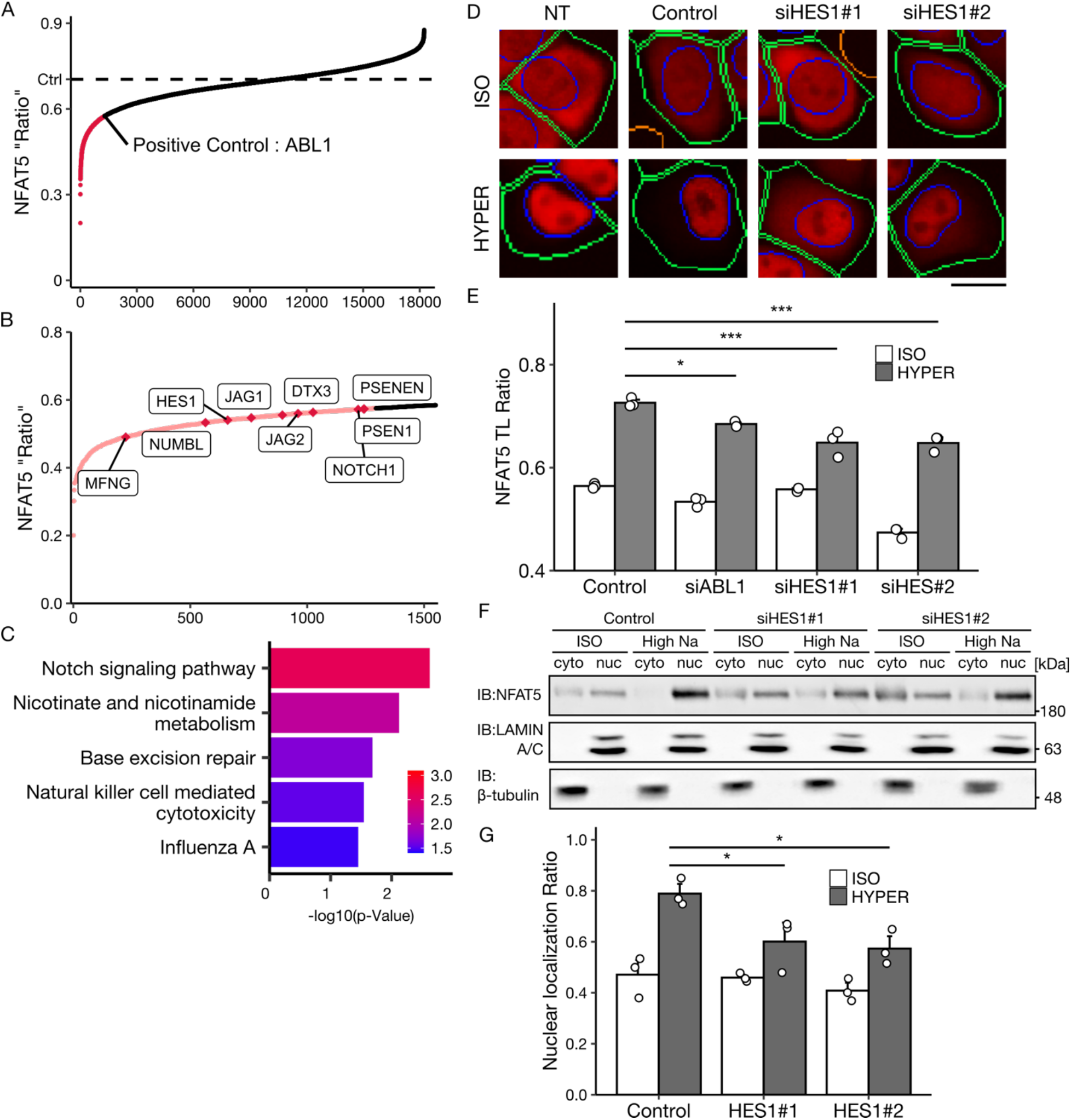
Figure 1. HES1 is necessary for NFAT5 nuclear translocation under high salt stress. (**A, B**) A genome-wide siRNA screen to search for regulators of NFAT5 nuclear translocation under high salinity/hyperosmotic stress, 500 mOsm; 3 h. (B) Notch-related genes in the positive hits whose knockdown attenuated NFAT5 nuclear translocation more than c-Abl1 knockdown. The vertical line shows the NFAT5 TL ratio, and the horizontal line shows the rank of genes. (**C**) KEGG pathway analysis of the 1291 positive hits. (**D, E**) Effect of HES1 depletion on NFAT5 nuclear translocation. Representative images of NFAT5-ΔtdTomato in the HeLa cells treated with siRNAs. Blue and green lines represent the borders of the nuclear and whole-cell regions, respectively. The scale bar represents 10 µm. NT: Nontransfected control. (E) NFAT5 TL ratio calculated from the fluorescent images (E, n = 3, analyzing 400-800 cells in each experiment). (**F, G**) Effect of HES1 depletion on the subcellular distribution of endogenous NFAT5 in HeLa cells. (F) LAMIN A/C and β-tubulin are nuclear and cytoplasmic marker proteins, respectively. NFAT5 TL ratio calculated from the band intensity (G, n = 5). In the bar graphs, individual values (white points) and the mean ± SEM are presented. *p < 0.05, ***p < 0.001. ISO, 300 mOsm; high Na, 500 mOsm in (D, E) and 400 mOsm in (F, G); 3 h. See also Fig. S1.

We validated the effect of the depletion of several Notch-related hit genes (HES1, PSEN 1/2 and Notch1) on NFAT5 nuclear translocation with individual siRNAs. Among the genes, knockdown of HES1 (Fig. S1C) most strongly attenuated NFAT5 nuclear translocation under high salt conditions (Fig. S1E). Two independent siRNAs against HES1 resulted in a similar effect on NFAT5 translocation (Fig. 1, D and E). In contrast, knockdown of Notch1 only slightly suppressed NFAT5 nuclear translocation, and knockdown of PSEN 1/2 showed almost no effect on the translocation (Fig. S1E). Thus, we hereafter focused on HES1 in NFAT5 regulation. To confirm the requirement of HES1 for endogenous NFAT5 nuclear translocation, we examined the subcellular distribution of endogenous NFAT5 by immunoblot analysis after fractionation (Fig. 1, F and G). In the control cells, endogenous NFAT5 protein was detected in both the cytoplasmic and nuclear fractions under isoosmotic conditions (TL ratio ≈ 0.53). Whereas the NFAT5 protein was mainly detected in the nuclear fraction after three hours under high salt conditions (TL ratio ≈ 0.81). In the HES1 knockdown cells, the NFAT5 protein was still observed in the cytoplasmic fraction under high salt conditions, and the TL ratio was significantly decreased compared with that of the control cells. A similar result was obtained in the immunocytochemical analysis of endogenous NFAT5 (Fig. S1, F and G). In contrast, HES1 depletion did not show any effect on Ca^2+^ ionophore-induced nuclear translocation of another NFAT family, NFATc3 (Fig. S1, H and I). These results suggested that HES1 is specifically required for high salt stress-induced NFAT5 nuclear translocation.

### HES1 is induced by ERK signaling in hyperosmotic stress and enhances NFAT5 nuclear translocation

Interestingly, we found that the HES1 protein amount was upregulated by hyperosmotic stress (Fig. 2A). HES1 was mainly localized in the nucleus under both iso- and high-salinity conditions (Fig. S2A). We then investigated the molecular mechanism of HES1 induction under high salt conditions. In addition to immunoblot analysis, RT-qPCR analysis suggested that the induction of HES1 is attributed to transcriptional regulation (Fig. 2B). HES1 is known to be induced by the Notch signaling pathway (*17*). Therefore, we first investigated the involvement of the canonical Notch signaling pathway in high salt-dependent HES1 induction. Treatment with DAPT, an inhibitor of gamma-secretase, which is a critical protease in the activation of the canonical Notch pathway (*18*), did not suppress high salt-dependent HES1 induction at either the mRNA or protein amount (Fig. S2, B and C). HES1 was also reported to be induced by gamma-secretase-independent noncanonical Notch signaling pathways (*19*). We focused on one of the noncanonical pathways, the ERK pathway, which was reported to increase the expression of HES1 in growth factor stimulation (*20*) because ERK activation in hyperosmotic stress has been reported (*21*). Consistent with previous studies, high salinity/hyperosmotic conditions enhanced the phosphorylation of ERK, which was completely suppressed by the MEK1/2 inhibitor U0126 (Fig. 2C). U0126 treatment attenuated the induction of HES1 under high salt stress at both the protein and mRNA amounts (Fig. 2, C and D). These results suggest that high salt stress promotes HES1 expression in an ERK pathway-dependent manner. Exogenous HES1 overexpression promoted the nuclear translocation of NFAT5Δ even under isosmotic conditions, suggesting that HES1 induction by high salt stress may further enhance NFAT5 translocation (Fig. 2E). The TL ratio quantified from the images of the HES1-overexpressing cells was significantly higher than that of the control cells under both iso- and high-salt conditions (Fig.2F). Consistent with the knockdown experiment, HES1 overexpression did not affect the localization of NFATc3 (Fig. S2, D and E), confirming the specificity to NFAT5.

**Fig. 2.**
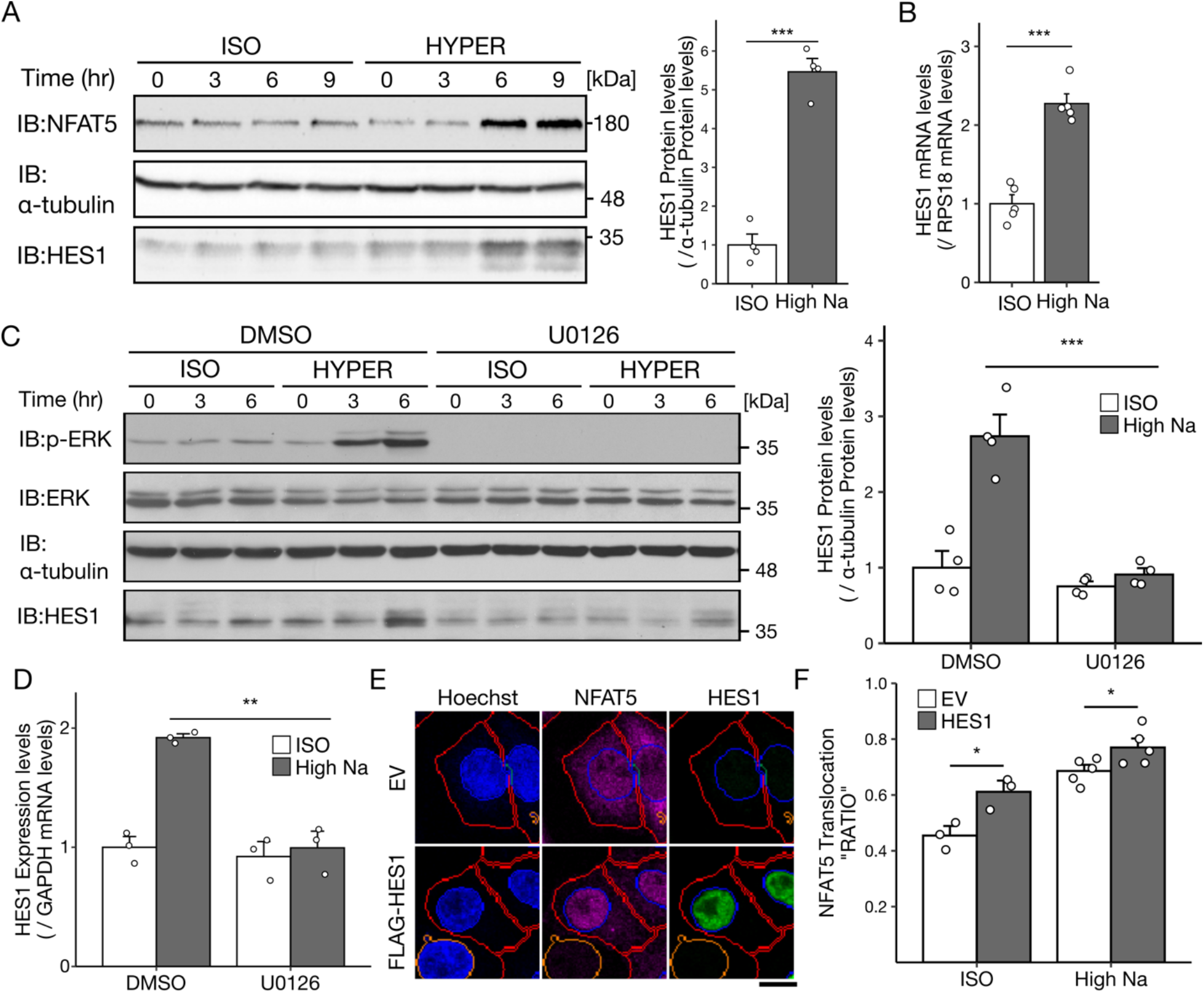
HES1 is induced by high salt stress via ERK signaling. (**A**) Time course of HES1 protein expression under high salt conditions in HeLa cells. The bar graph indicates HES1 protein amount at 6 h after osmotic stimuli (n = 4). (**B**) mRNA expression of Hes1 under high salt conditions in HeLa cells (n = 5). mRNA expression in each sample was normalized to the mean expression of samples under isoosmotic conditions. (**C, D**) Effect of the ERK signaling inhibitor U0126 on high salt stress-induced HES1 protein. The bar graph indicates HES1 protein amount at 6 h after osmotic stimuli (C, n = 3 and mRNA expression (D, n = 3). Cells were pretreated with 5 µM U0126 for 30 min. DMSO: dimethyl sulfoxide. mRNA expression in each sample was normalized to the mean expression of samples with DMSO treatment under isoosmotic conditions. (**E, F**) Effect of exogenous HES1 expression on NFAT5 nuclear translocation. Representative images of NFAT5Δ-tdTomato in HeLa cells under isoomolarity. The scale bar represents 10 µm. EV: empty vector, a negative transfection control (E). NFAT5 TL ratio calculated from the fluorescent images (F, n = 3-5, analyzing 40-80 cells in each experiment). In the bar graphs, individual values (white points) and the mean ± SEM are presented. *p < 0.05, **p < 0.01, ***p < 0.001. ISO, 300 mOsm; high Na, 400 mOsm; 5 h in (B, D) and 3 h in (F). See also Fig. S2.

### HES1 is required for induction of osmoprotective genes that are regulated by NFAT5

The nuclear translocation of NFAT5 under hyperosmotic conditions is known to be important for the induction of osmoprotective genes such as chaperons, osmolyte synthases and transporters (*4*). We next examined whether HES1 affects NFAT5-mediated gene induction under high salinity/hyperosmotic conditions (Fig. 3). We confirmed by RT-qPCR that high salt stress markedly increased the expression of the representative NFAT5 target genes betaine/GABA transporter 1 (BGT1) and aldose reductase (AR) (Fig. 3, A and B). The induction was clearly abrogated by NFAT5 depletion. HES1 knockdown also significantly attenuated BGT1 expression under high salt conditions (Fig. 3A). However, surprisingly, the induction of AR was not affected in the HES1 knockdown cells (Fig. 3B). These results suggested that HES1 may be involved in the regulation of a portion of NFAT5 target genes.

**Fig. 3.**
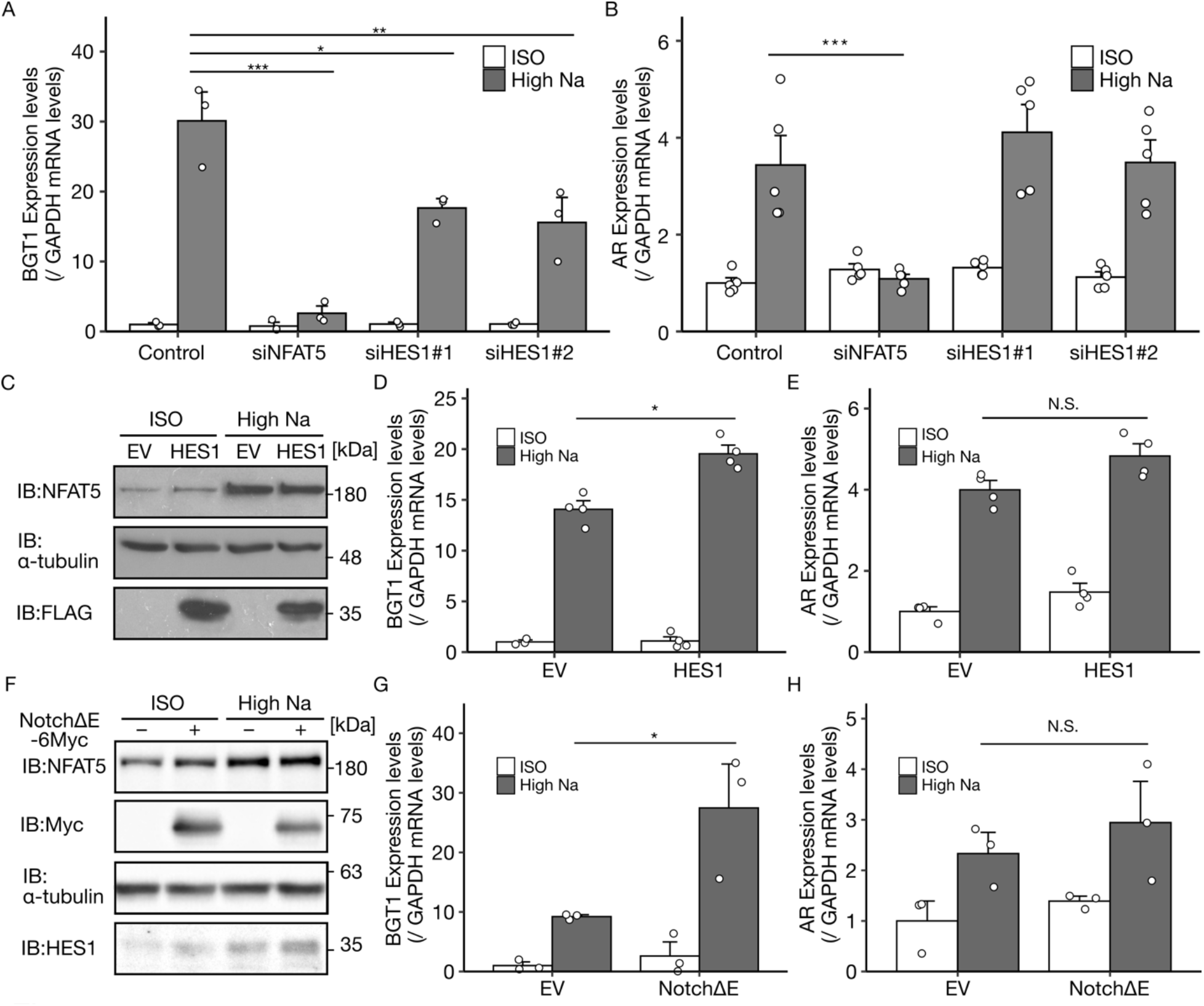
HES1 regulates the induction of a NFAT5 target gene, BGT1, but not that of another one, Aldose Reductase under high salt stress. (**A, B**) Effect of HES1 depletion on the NFAT5 target genes BGT1 (A, n = 3) and AR (B, n = 5). mRNA expression in each sample was normalized to the mean expression of samples with control siRNA under isoosmotic conditions. (**C**) Expression of HES1 and NFAT5 in the HeLa cells transiently transfected with the EV or FLAG-HES1 construct. (**D, E**) Effect of exogenous HES1 expression on the NFAT5 target genes BGT1 (D, n = 4) and AR (E, n = 4). (**F**) Endogenous HES1 expression in the HeLa cells transfected with a constitutively active Notch1 (NotchΔE-6Myc) construct. (**G, H**) Effect of exogenous NotchΔE expression on the NFAT5 target genes BGT1 (G, n = 3) and AR (H, n = 3). In the bar graphs, individual values (white points) and the mean ± SEM are presented. N.S., not significant; *p < 0.05, **p < 0.01, ***p < 0.001. ISO, 300 mOsm; high Na, 400 mOsm; 7.5 h. See also Fig. S3.

We next examined the effect of HES1 overexpression on NFAT5-mediated gene induction under high salt conditions. Consistent with the knockdown experiment, HES1 overexpression significantly increased BGT1 induction but not AR induction (Fig. 3, C to E). These results suggested that HES1 upregulation leads to further induction of restricted NFAT5 target gene(s) under high salt conditions. In support of this idea, the expression amounts of BGT1, a HES1-regulated NFAT5 downstream gene (Fig. 3A), were markedly induced at a relatively late time point, 7.5 hours (when HES1 was upregulated) after high salt stimuli (Fig. S3A). In addition, the attenuation of BGT1 induction by HES1 knockdown was more clearly observed at 7.5 hours than at 5 hours (Fig. S3B).

We further investigated the effect of constitutively active Notch1 (NotchΔE) (*22*) on NFAT5-dependent gene induction. Transfection of NotchΔE enhanced HES1 protein expression (Fig. 3F), since HES1 is a target of canonical Notch signaling (*23*). In this situation, as well as HES1 overexpression, NotchΔE transfection increased the expression of BGT1 transcripts relative to the control (Fig. 3G) but not those of AR (Fig. 3H). These results demonstrated that HES1 induction via canonical Notch pathway signaling can also promote NFAT5 activity on restricted gene(s) in high salt conditions, suggesting that costimulation of the Notch signaling pathway can modulate the cellular high salinity/hyperosmotic response through HES1 induction.

To comprehensively identify which NFAT5 target genes were regulated by HES1, we performed a microarray analysis (Fig. 4A). Among 457 genes that showed induction (fold > 2.0) upon high salt stress in the control cells, 132 genes were NFAT5 dependent, as NFAT5 knockdown suppressed the induction (NFAT5 KD/control < 0.5 under high salt stress) (Fig. 4B). HES1 knockdown also suppressed the induction of 81 genes (61.4%) among the 132 NFAT5-dependent genes (HES1 KD/control < 0.67 under high salt stress). BGT1 was included in the category (both NFAT5- and HES1-dependent), and AR was excluded, confirming previous RT-qPCR results (Fig. 3, A and B). A KEGG pathway analysis of the 81 common genes revealed that genes involved in amino acid and sugar metabolism were enriched in both NFAT5- and HES1-dependent genes (Fig. 4C), e.g., ARG2, GLS and SLC2A1. We confirmed the dependence of these genes on both NFAT5 and HES1 by RT-qPCR (Fig. 4, D to F and Fig. S3, C to E). Several amino acids and sugars are known as osmolytes that are important for cell survival in hyperosmotic conditions (*2, 24*). Thus, these results suggest that HES1 may have a cytoprotective function in high salinity/hyperosmotic conditions by promoting a considerable portion of NFAT5 target gene induction.

**Fig. 4.**
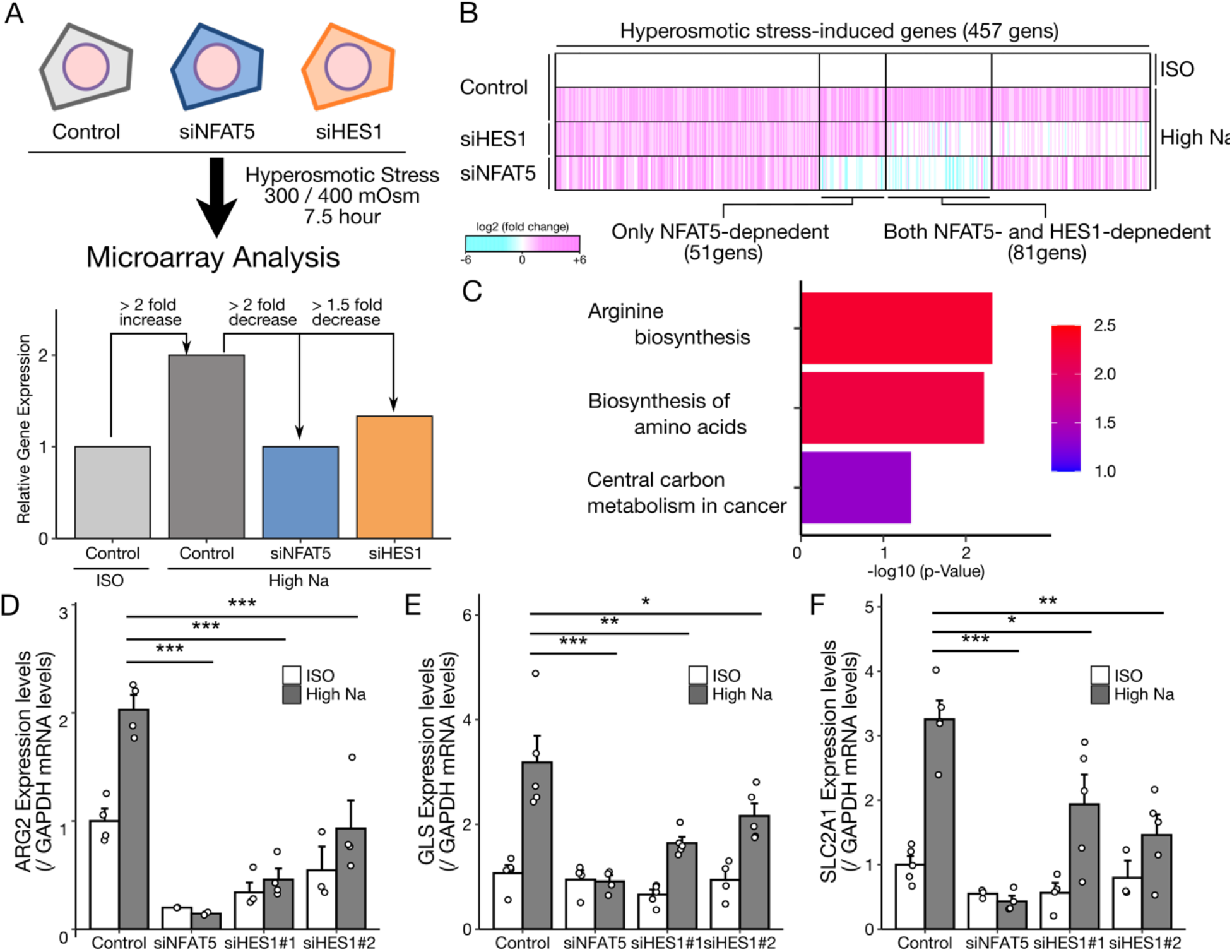
HES1 promotes the induction of multiple NFAT5 target genes under high salt stress. (**A**) Schematic model showing the design of the microarray analysis and gene classification. (**B**) Heatmap showing the log fold change in the mRNA expression of high salt stress-induced genes. The expression under high salt (400 mOsm) with each siRNA was compared with that under isoosmolarity (300 mOsm) with control siRNA. (**C**) KEGG pathway analysis of both NFAT5- and HES1-dependent genes (81 genes) in the microarray analysis. (**D-F**) Effect of HES1 depletion on both NFAT5- and HES1-dependent genes identified by microarray analysis, ARG2 (D, n = 4), GLS (E, n = 5) and SLC2A1 (F, n = 5). In the bar graphs, individual values (white points) and the mean ± SEM are presented. N.S., not significant; *p < 0.05, **p < 0.01, ***p < 0.001. ISO, 300 mOsm; high Na, 400 mOsm; 7.5 h. See also Fig. S3.

### HES1 functions in cytoprotection under hyperosmotic stress

As NFAT5 plays a pivotal role in cell survival under hyperosmotic conditions, we investigated the involvement of HES1 in the protection against cell death induced by chronic hyperosmotic stress. Similar to NFAT5 knockdown, HES1 knockdown enhanced the number of propidium iodide (PI)-positive cells and LDH release under high salt conditions, suggesting aggravation of necrotic cell death (Fig. 5, A and B). Moreover, the activity of an apoptotic marker, caspase 3, was significantly elevated in either the NFAT5- or HES1-depleted cells compared with the control cells under hyperosmotic conditions (Fig. 5C). Additional depletion of HES1 did not accelerate caspase 3 activity in the NFAT5-deficient cells (Fig. S3F). Furthermore, the exogenous overexpression of BGT1 partially but significantly suppressed the induction of apoptosis in either the HES1- or NFAT5-depleted cells (Fig. S3G). Collectively, these results suggest that HES1 is important for cell survival under high salinity/hyperosmotic conditions via regulation of NFAT5-mediated gene expression.

**Fig. 5.**
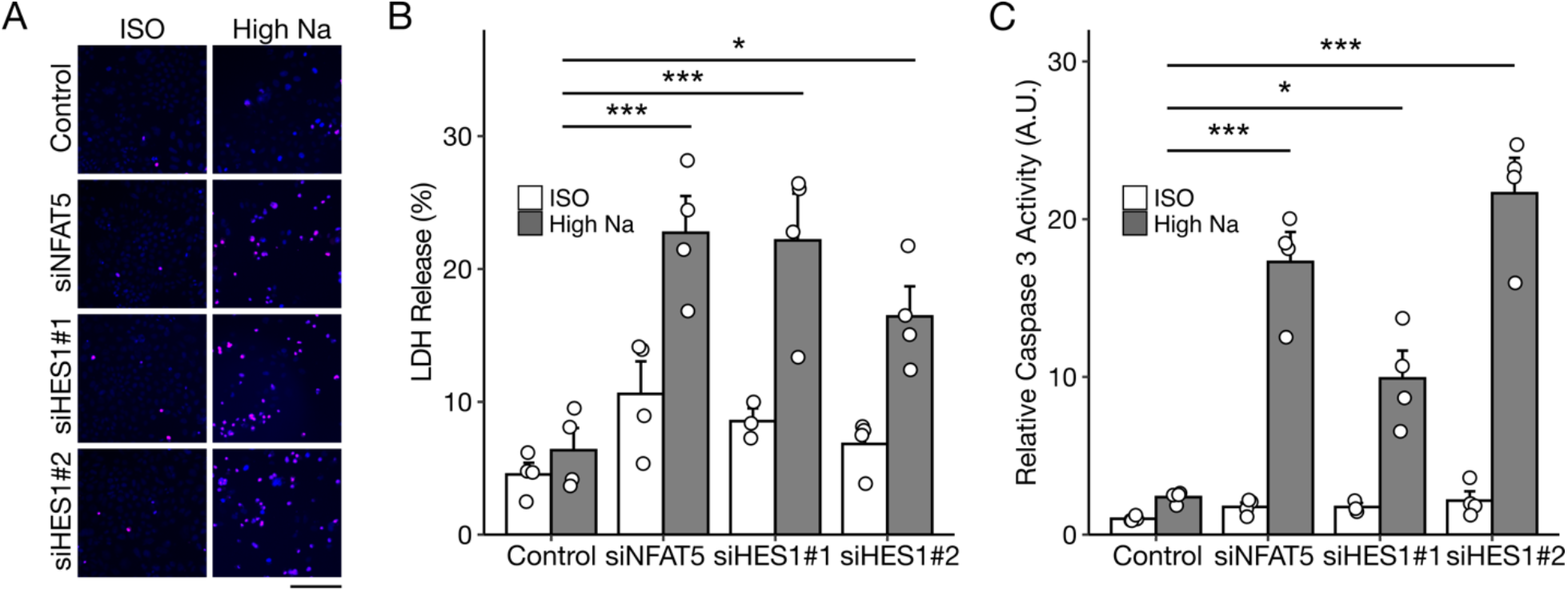
HES1 has cytoprotective functions against high salinity/hyperosmotic stress. (**A-C**) Effect of HES1 depletion on high salt stress-induced cell death. (A) Propidium iodide, PI staining (representative data, n = 3), lactate dehydrogenase (LDH) release assay (B, n= 4) and caspase 3 activity (C, n = 4). Caspase 3 activity in each sample was normalized to the mean activity of samples with control siRNA under isoosmotic conditions. In the bar graphs, individual values (white points) and the mean ± SEM are presented. N.S., not significant; *p < 0.05, **p < 0.01, ***p < 0.001. ISO, 300 mOsm; high Na, 400 mOsm; 24 h in (A), 15 h in (B) and 9 h in (C). The scale bar represents 100 µm. See also Fig. S3.

### HES1 regulates NFAT5 activity through the recruitment of NFAT5 to the target promoter

To determine the molecular mechanism by which HES1 promotes NFAT5 target gene induction, we investigated NFAT5 recruitment to the promoter region of its target gene. We found two potential NFAT5 binding motifs (TGGAAANNYNY) within −2 kbp of the transcription start site (TSS) in the two representative genes, BGT1 (both NFAT5- and HES1-dependent) and AR (NFAT5-dependent and HES1-independent) (Fig. 6, A and B). Consistent with previous studies (*25, 26*), NFAT5 recruitment to these binding motifs ((i) and (ii) in each gene) was elevated by high salt stress in the control cells (Fig. 6, C to F). NFAT5 knockdown diminished these signals, confirming the specificity of the antibody used in the ChIP-qPCR analysis. High salt stress-induced NFAT5 recruitment was significantly suppressed by HES1 knockdown at the two binding motifs in the BGT1 promoter (Fig. 6, C and D). However, HES1 knockdown did not suppress NFAT5 recruitment to the two binding motifs in the AR promoter (Fig. 6, E and F). These results suggest that HES1 may specifically enhance the recruitment of NFAT5 to the promoter of HES1-dependent NFAT5 target genes under high salt stress.

Although it is well known HES1 binding to promoters results in transcriptional repression (*27*), some studies reported that HES1 can function as a transcriptional activator at the promoter of certain genes (*28, 29*). Therefore, we speculated that HES1 might bind to the promoter region of BGT1 to promote gene expression. A database of transcription factor-binding profiles, JASPAR (*30*), predicted 5 candidate sites of the HES1 binding motif (a-e) within the −2 kbp region of the BGT1 TSS (Fig. 6G). Among them, we focused on site c, which is adjacent to one of the NFAT5 binding motifs (Bgt1-(ii)) (Fig. 6G). ChIP-qPCR analysis with an anti-HES1 antibody revealed that HES1 recruitment around site c was significantly elevated by high salt stress, which was abrogated by HES1 knockdown (Fig. 6H). This result suggested that high salt stress induces HES1 recruitment to the adjacent region of the NFAT5 binding site in the BGT1 promoter.

**Fig. 6.**
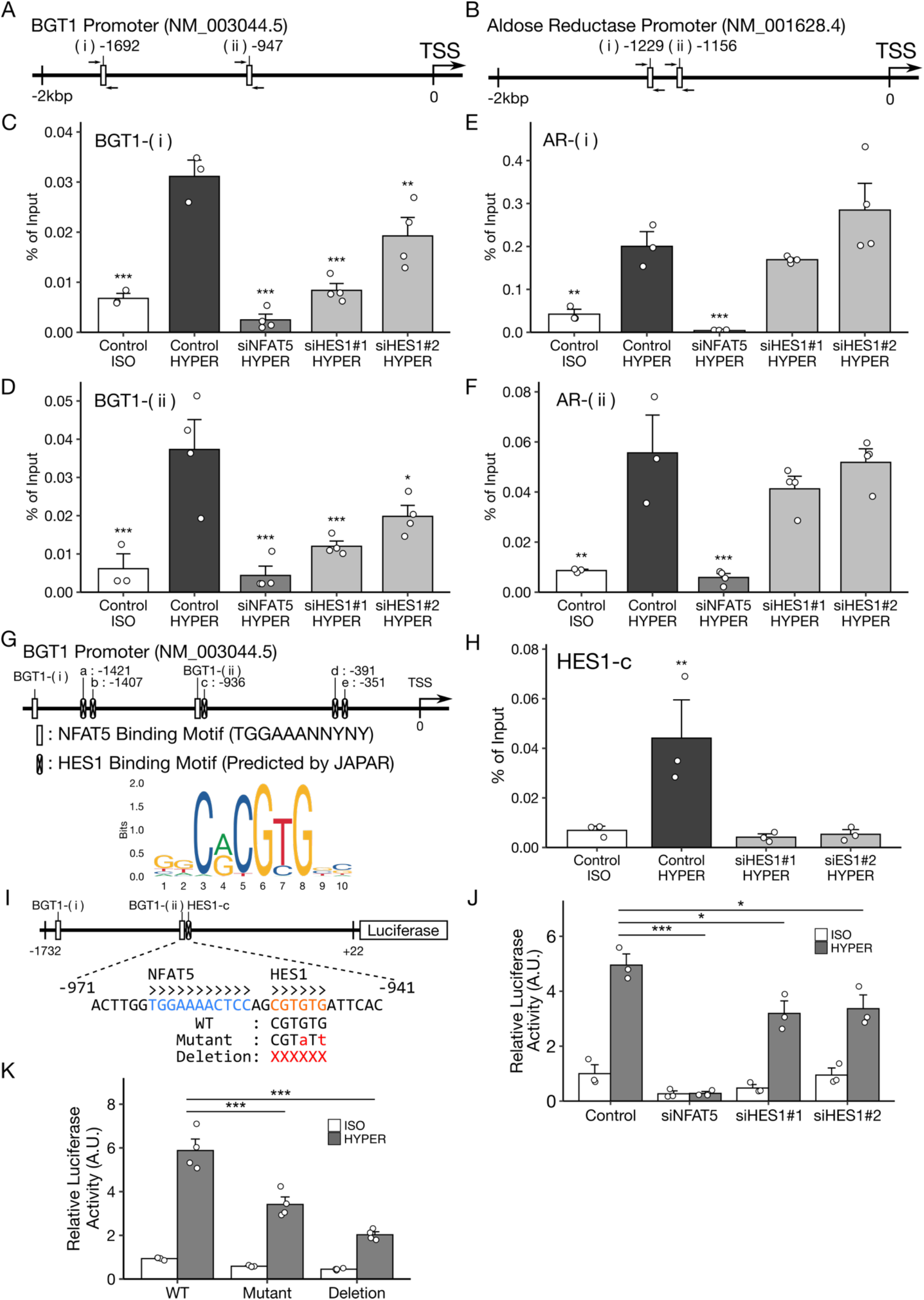
HES1 regulates BGT1 promoter activity through the recruitment of NFAT5 to the target motif. (**A, B**) Schematic models showing the promoter regions of the NFAT5 target genes BGT1 (A) and AR (B). Each has two NFAT5-binding motifs, (i) and (ii), presented as white boxes. (**C-F**) ChIP analysis investigating the effect of HES1 depletion on the DNA binding of NFAT5 to each motif, BGT1-(i) and -(ii) and AR-(i) and -(ii) (n = 4). (**G**) A schematic model showing the prediction of HES1 binding motifs in the BGT1 promoter using JASPAR. The predicted sites of HES1 binding are presented as white boxes with x-markers. (**H**) ChIP analysis evaluating HES1 recruitment to a predicted site (HES1-c site) in the BGT1 promoter under high salt conditions. (**I**) A schematic model showing the structure of the BGT1 reporter vector and mutations of the HES1 binding site introduced into the reporter. (**J**) Effect of HES1 depletion on BGT1 reporter activity (n = 3). (**K**) Effect of a double point mutant (Mutant) and deletion mutant (Deletion) in the HES1 binding site on the BGT1 reporter activity (n = 4). Mutant sequences shown in (I). In the bar graphs, individual values (white points) and the mean ± SEM are presented. *p < 0.05, **p < 0.01, ***p < 0.001. ISO, 300 mOsm; high Na, 400 mOsm; 7.5 h in (C-F and J), 5.5 h in (H) and 9 h in (I). See also Fig. S4.

To investigate the functional role of the HES1 binding motif in the BGT1 promoter, we constructed a luciferase reporter that contains the −1732 to 22 bp region of the BGT1 promoter (Fig. 6I). In the luciferase reporter assay, reporter activity was significantly increased upon high salt stress, and it was completely and moderately suppressed by NFAT5 and HES1 knockdown (Fig. 6J), respectively, confirming that the reporter exhibits similar properties to endogenous BGT1 gene expression (Fig. 3A). In addition, by mutating the NFAT5 binding motifs, we confirmed that the two NFAT5 binding motifs in the construct were functional for reporter activity (Fig. S4A). Then, we introduced mutations in HES1 binding motif-c of the BGT1 reporter construct (Fig. 6I). Both base substitutions and complete deletion of 6 bases in the HES1 binding motif significantly attenuated reporter activity under high salt conditions (Fig. 6K).These results suggest that an HES1 binding motif adjacent to the NFAT5 binding site in the BGT1 promoter is required for BGT1 induction under high-salt conditions. In parallel, we found that the HES1 mutant lacking the DNA binding ability (ΔbHLH) (*27*) could not enhance NFAT5 nuclear translocation or BGT1 expression (Fig. S4, B to D). Collectively, our results suggest that HES1 binding to the promoter region enhances NFAT5 recruitment and thus the transcription of NFAT5 target genes under high-salt conditions.

## DISCUSSION

In this study, through a genome-wide siRNA screen, we found that HES1 is necessary for the nuclear translocation of NFAT5 (Fig. 1 and Fig. 2) and its recruitment to the DNA promoter under high salinity/hyperosmotic conditions (Fig. 6). Although HES1 is known to function as a transcriptional repressor, in the present context, HES1 activates the transcription of osmo-protective genes and contributes to cellular tolerance against high salinity/hyperosmotic stress (Fig. 3 to 5).

Regarding the mechanism by which HES1 promotes NFAT5 nuclear translocation, HES1 exhibits specificity to NFAT5, as the translocation of another NFAT family, NFATc3, was not affected by HES1 depletion and overexpression (Fig. 1, D and E, Fig. S1, H and I, Fig. 2, E and F, Fig. S2, D and E). We speculate that HES1 may promote NFAT5 nuclear translocation by recruiting NFAT5 to the target promoter region (Fig. 6) because it has been reported that DNA binding determines NFAT5 nuclear translocation (*31*). The promoter analysis of BGT1 suggested that a HES1 binding region is located in close proximity to an NFAT5 binding motif and is important for NFAT5 recruitment (Fig. 6, G to K). In addition, bioinformatics analysis using JASPAR revealed that several promoter regions of both NFAT5- and HES1-dependent genes (e.g., SLC2A1 and LULAP1L) possess HES1 binding motifs in the vicinity of the NFAT5 motif (within 20 bases). HES1 binding motifs in the AR promoter, one of the only NFAT5-dependent genes, are over 700 bp apart from NFAT5 binding motifs. Thus, the adjacent HES1 binding motif might be important to recruit NFAT5 to its binding motif. NFAT5 recruitment may be mediated by the direct interaction between HES1 and NFAT5. HES1 induced by high salt stress may function as a scaffold molecule to promote NFAT5 recruitment to adjacent binding regions. Another possibility is regulation through chromatin remodeling by HES1, as HES1 promotes the recruitment of chromatin-modifying enzymes (*32*). Some studies have reported that high salinity dynamically changes chromatin modifications (*33*), especially in BGT1 (*34*). HES1 may be involved in such remodeling of chromatin structure, thereby recruiting NFAT5 to the adjacent DNA binding motifs. Further studies of the molecular mechanism of HES1 as a transcriptional activator for selective NFAT5 target genes under high salt conditions may reveal a novel function of this well-studied Notch-related transcription factor.Here, we attempted to comprehensively search for molecules regulating NFAT5 nuclear translocation through a genome-wide siRNA screen. The core molecules of nuclear transport and several known NFAT5 regulators were highly ranked (e.g., KPNB1, RANBP2, CDK5 and ABL1, Data file S1), suggesting that the screen was conducted successfully. In this study, we focused on HES1, which is involved in Notch signaling and exhibited the highest score in the pathway analysis.

HES1 induction through Notch signaling is known to play key roles in cell differentiation and developmental processes (*16*). However, to date, little is known about the relationship between Notch signaling and the hyperosmotic stress response. In *Caenorhabditis elegans*, nonsense mutants of Notch ligands, OSM11 and OSM7, fail to avoid high osmolarity (*35*). This report may suggest the involvement of Notch signaling in the hyperosmotic response, but these mutants show resistance to stress via accumulation of glycerol as an osmolyte. Although the contribution to the response might be in the opposite direction, our finding expands the role of Notch signaling in the hyperosmotic environment in mammalian cells with a new molecular interaction between HES1 and NFAT5.

Although other Notch-related genes were included among the positive genes (Fig. 1B), their repression with individual siRNAs had only a limited effect on NFAT5 (Fig. S1E). We speculated that Notch signaling might be activated for some reasons and that HES1 was elevated in the screening condition; thus, the contribution of Notch signaling to NFAT5 regulation might be increased. Supporting our inference, forced activation of Notch signaling by a constitutively active mutant led to HES1 induction and enhanced the expression of BGT1 (Fig. 3, F and G). Therefore, Notch-related genes other than HES1 might also affect NFAT5 nuclear translocation when Notch signaling is activated.

In addition, the screen provides new insights into osmosensing in mammalian cells. Nicotinate and nicotinamide metabolism also exhibited high scores in the pathway analysis in our screen (Fig. 1C). Recently, we reported that poly(ADP-ribose) and nicotinamide metabolic enzymes modulate the acute hyperosmotic stress response through the regulation of an osmoresponsive kinase (ASK3) (*36*). A common mechanism through nicotinamide metabolism might regulate osmosensing in both acute (kinase regulation) and chronic (NFAT5-mediated gene induction) hyperosmotic responses.

We also found that high salt stress enhanced HES1 expression via a noncanonical Notch signaling, ERK signaling. In cancer cells, ERK signaling is excessively activated and is a potent target for therapy (*37*). In renal carcinoma cells, constitutive activation of ERK1/2 enhances cell proliferation and migration through NFAT5 activation even under isoosmotic conditions (*38*). Similar oncogenic phenotypes induced by NFAT5-mediated glycolytic gene expression are observed in pancreatic cancer (*39*). Aberrant ERK signaling might enhance the HES1-NFAT5 axis, thereby promoting cancer progression.

In this study, we found that HES1 protects against high salt stress-induced cell death (Fig. 5, A to C). Considering that cell death under high salinity/hyperosmotic stress is caused by various mechanisms (e.g., apoptosis and necrosis) (*2*), it is likely that HES1 is involved in the global regulation of vulnerability to stress rather than a specific signaling of cell death. Exogenous overexpression of BGT1 restored cellular tolerance to stress in the HES1 and NFAT5 knockdown cells, but the recovery was only partial (Fig. S3G). The microarray analysis revealed that HES1 broadly influences NFAT5-mediated gene induction related to osmolytes such as SLC2A1, GLS, ARG2 and BGT1 (Fig. 3 and 4). The pleiotropic regulation of osmolytes by HES1 through NFAT5 regulation might be necessary for cytoprotection against high salinity/hyperosmotic stress.

Recently, NFAT5 has attracted interest for immune cell regulation in high-salt environments (*5*). Activation of NFAT5 by high salinity induces proinflammatory cell polarization and cytokine production in monocytes and T cells. The NFAT5-mediated inflammatory response not only enhances antitumor activity (*7*) and antimicrobial activity (*8*) but also leads to the development and exacerbation of immune-related diseases such as atopic dermatitis (*10*) and experimental autoimmune encephalomyelitis (EAE) (*40*). Interestingly, lymphoid tissues such as the thymus and spleen were reported to show higher osmolarity than other tissues and plasma (*41*). The thymus is a primary lymphoid tissue for T-cell development in which Notch signaling, including HES1, is essential (*42*). Although the precise role of hyperosmolarity in lymphoid tissues awaits further research, the present HES1 function in high salinity/hyperosmotic conditions may regulate immune cell maturation through NFAT5 regulation. In a rare example of HES1 as a transcriptional activator, VEGF-C expression in bone marrow-derived macrophages (BMDMs) is promoted by HES1 binding to its promoter (*28*). Intriguingly, VEGF-C is one of the NFAT5 target genes in macrophages (*43*), and it is possible that NFAT5 and HES1 cooperatively modulate VEGF-C expression and subsequent inflammation. Immune regulation through the HES1-NFAT5 axis would be an attractive topic for future studies. NFAT5 was originally identified as an osmoresponsive factor but is now known to respond to a broad range of stresses such as hyperlipidemia, ischemia and inflammation (*5*). The fine-tuning mechanisms that determine the induction of different genes through NFAT5 depending on the stresses are still unclear. Specific interactors of NFAT5 on the promoters of target genes might be important for regulation under individual conditions, e.g., DNMT1 in hyperlipidemia (*44*) and NFκB in inflammation (*45*). In addition to these molecules, HES1 might shape the specific character of NFAT5 under high salinity/hyperosmotic stress.

## MATERIALS AND METHODS

### Cell lines and cell culture

HeLa cells were cultured in Dulbecco’s Modified Eagle’s Medium (DMEM)-low glucose (Sigma-Aldrich, Cat#D6046) supplemented with 10% fetal bovine serum (FBS; BioWest, Cat#S1560-500) and 100 units/mL penicillin G (Meiji Seika, Cat#6111400D2039). HeLa cells stably expressing NFAT5Δ547-tdTomato (NFAT5Δ) and GFP were established by transfecting with NFAT5Δ in pcDNA6 vector (Invitrogen) and EGFP in pcDNA4/TO vector (Invitrogen). Cells were selected with blasticidin (Invitrogen, Cat. #A1113903) and Zeocin (Invitrogen, Cat. #R25001), then picked for single clones, and cultured in DMEM-low glucose supplemented with 10% FBS, 2.5 μg/mL blasticidin and 50 μg/mL Zeocin. All cells were cultured in 5% CO2 at 37°C and verified to be negative for mycoplasma.

### Transfections

Plasmid transfections were performed with polyethylenimine “MAX” (Polysciences, Cat#24765) to cells at 70% confluency, according to a previously described protocol (Longo et al., 2013) with minor optimization. siRNA transfections were carried out by reverse transfection using Lipofectamine RNAiMAX (Invitrogen, Cat. #133778-500) and 10 nM siRNAs (Horizon Disxovery, siGENOME siRNAs), according to the manufacturer’s instructions. In combination transfections of both plasmid and siRNA, siRNA transfections were performed 12–24 h before plasmid transfections.

### Expression plasmids

Expression plasmids for this study were constructed by standard molecular biology techniques and all constructs were verified by sequencing. A human NFAT5 cDNA (CDS of NM_006599.4) lacking C-terminus (NFAT5Δ) [contains aa 1-547] was cloned from a cDNA pool derived from HeLa cells and subcloned into pcDNA3/GW (Invitrogen) with a C-terminal tdTomato-tag. A human HES1 cDNA (CDS of NM_005524.4) was subcloned into pcDNA3/GW (Invitrogen) with an N-terminal Flag-tag from EF.hHES1.Ubc.GFP, gifted from Linzhao Cheng (Addgene plasmid # 17624; http://n2t.net/addgene:17624; RRID:Addgene_17624). A mouse Notch1 lacking the extracellular region (Notch1ΔE) [contains aa 1-23 and then aa 1704-2184] with C-terminal 6-Myc tag (Hofelmayr et al., 1999) was subcloned into pcDNA3/GW from pCS2 Notch1 ΔEMV-6MT, kindly gifted from Dr. Raphael Kopan (Addgene plasmid #41737; http://n2t.net/addgene:41737; RRID:Addgene_41737). A human NFATc3 cDNA (CDS of NM_004555.4) was cloned from a cDNA pool derived from HeLa cells and subcloned into pcDNA3/GW with a N-terminal GFP-tag.

### Osmotic stress treatments

High salinity / hyperosmotic stress was treated by applying 2.25 M NaCl solution into culture medium (≈ 300 mOsm/kg H2O) to add 50 mM or 100 mM NaCl, corresponding with 400 or 500 mOsm/kg H2O, respectively. Isoosmotic stress was applied by adding the same volume of 150 mM NaCl solution. Absolute osmolality was verified by Osmomat 030 (Gonotec) to fall within a range of ± 25 mOsm /kg H2O.

### Cell lysis and western blotting

Cells were lysed in a lysis buffer (20 mM Tris-HCl pH 7.5, 150 mM NaCl, 10 mM EDTA, 1% sodium deoxycholate, and 1% Triton X-100) supplemented with 1 mM phenylmethylsulfonyl fluoride (PMSF), 5 μg/mL leupeptin. Cell extracts were centrifuged and supernatants were sampled by adding 2x SDS sample buffer (80 mM Tris-HCl pH 8.8, 80 μg/mL bromophenol blue, 28.8% glycerol, 4% SDS and 10 mM dithiothreitol). After boiling at 98℃ for 3 min, the samples were resolved by SDS-PAGE and electroblotted onto a BioTrace PVDF membrane (Pall), FluoroTrans W membrane (Pall) or Immobilon-P membrane (Millipore, Cat#IPVH00010). The membranes were blocked with 5% skim milk (Megmilk Snow Brand) in TBS-T (20 mM Tris-HCl pH 8.0, 137 mM NaCl and 0.1% Tween 20) and probed with the appropriate primary antibodies diluted by 1st antibody-dilution buffer (TBS-T supplemented with 5% BSA (Iwai Chemicals, Cat#A001) and 0.1% NaN3 (Nacalai Tesque, Cat#312-33)). After replacing and probing the appropriate secondary antibodies diluted by skim milk in TBS-T, antibody-antigen complexes were detected on X-ray films (FUJIFILM, Cat#47410-07523, Cat#47410-26615 or Cat#47410-07595) using an ECL system (GE Healthcare). Quantification was performed via densitometry using Fiji/ImageJ software (Schindelin et al., 2012). “TL ratio” were defined as the ratio of protein detected in nuclear sample to sum of that in nuclear and cytosolic sample.

### Subcellular fractionation

For preparation of subcellular fraction, cells on a culture plate were gently soaked in a cytoplasmic lysis buffer (10 mM HEPES, 15 mM MgCl2⋅6H2O1, 10 mM KCl, 0.5 v/v% NP-40) supplemented with 1 mM PMSF, 5 μg/mL leupeptin. After incubation at 4℃, lysates in the plate were centrifuged and supernatants were sampled by adding 2x SDS sample buffer as a cytoplasmic protein sample. After gently washing the plate once with the cytoplasmic lysis buffer, cells were lysed with in the lysis buffer with PMSF and leupeptin. Cell extracts were centrifuged and supernatants were sampled by adding 2x SDS sample buffer as a nuclear protein sample. The samples were analyzed using the same protocol of western blotting as mentioned above.

### Nuclear translocation measurements

Cells were fixed with 3.7% paraformaldehyde (Wako, Cat#064-00406) in PBS and stained with Hoechst 33258 in PBS containing 1% Triton X-100 (Sigma-Aldrich, Cat#T9284). NFAT5 Nuclear Translocation were measured and analyzed by using CellInsight NXT (Thermo Fisher Scientific) with the optimized Cell Health Profiling BioApplication (Fig. S1A). NFAT5 nuclear translocation ratio in each cell was calculated as the ratio of NFAT5 in nuclear region defined by Hoechst staining to that in whole cell region defined by GFP. In analysis with HeLa WT cells, Circ Target Region identified with nucleus was defined as whole cell region by the application. “NFAT5 TL ratio” in each condition was the mean of the translocation ratio of all cells from nine fields in a well (around 1000 cells in total). GFP-NFATc3 translocation was measured in a similar way.

### High-throughput assay system for NFAT5 nuclear translocation

We developed a fluorescent image-based assay system for NFAT5 nuclear translocation. In the following semi-automated protocol, solution was added and respirated using a Multidrop Combi (Thermo Fisher Scientific) and a AquaMax 2000 (Molecula Devices), respectively.

For siRNA transfection, 375 nM siRNA in OPTI-MEM was spotted onto 384-well plates, and Lipofectamine RNAiMAX in OPTI-MEM (1:100, Invitrogen, Cat#31985) was added at 10 μL/well. After 15 min, 5.0 × 104 NFAT5Δ and GFP expressing HeLa cells /mL were seeded at 40 μL/well (2,000 cells/well) onto the siRNA complex; the final siRNA concentration was 30 nM. After 48 h, culture medium was aspirated to 20 μL/well and high salt stress was applied by adding 2x NaCl (200 mM) in culture medium (final ≈ 500 mOsm/kg H2O) at 20 μL/well. To apply isoosmotic stress to reference wells, isoosmotic culture medium was used instead of NaCl-supplied medium. After 3 hours of osmotic stress, cells underwent the following fixation and nuclear staining steps: aspiration to 10 μL/well, fixation for 10 min by 2x paraformaldehyde (Wako Pure Chemical Industries, Cat#064-00406) in PBS (7.4%) at 20 μL/well (final 3.7% formaldehyde), 2 washes with PBS, aspiration to 10 μL/well, 15 min permeabilization and nuclear staining by overlaying 2x Triton X-100 (Sigma, Cat#T9284) and Hoechst 33258 (Dojindo, Cat#343-07961) in PBS (0.4%) at 10 μL/well (final 0.2% Triton X-100), 2 washes with PBS, aspiration to 10 μL/well, adding PBS to 100 μL/well. Finalized assay plates were measured and analyzed using the CellInsight NXT with an optimized Cell Health Profiling BioApplication (Fig. S1A).

### siRNA libraries and assay plate preparation

The genome-wide siRNA library used in the primary screen was consisted of 3 Dharmacon siGENOME SMARTpool collections: Human Genome (Cat#G-005005-02), Human Drug Targets (Cat#G-004655-02) and Human Druggable Subsets (Cat#G-004675-02). The library targeted 18,099 human genes, and each siRNA reagent was allocated as a pool of 4 different siRNAs to each well in 384-well plate format. The supplied lyophilized siRNA reagents were dissolved at 2 μM in 1x siRNA buffer (Dharmacon, Cat#B-002000-UB-100) with the Multidrop Combi. The master plate was copied and diluted to 375 nM on the intermediate plate (ABgene, Cat#AB-0781) using a Biomek FXP (Beckman Coulter). Assay plates were created by spotting 4 μL/well (1.5 pmol/well) from the intermediate plate onto 384-Well Optical Bottom Plates (Nunc, Cat#152041, same lot in the screening) with the Biomek FXP. For quality control during the screen, 375 nM control siRNA (Dharmacon, Cat#D-001206-13), ABL1 siRNA (Dharmacon, Cat#M-00310-07) and 1x siRNA buffer (for siRNA(–) control) were also spotted onto columns #1, 2, 23 and 24 of the assay plates. The plates were sealed and frozen at −30℃ until the assay was performed.

### Genome-wide siRNA screen

Using the established assay system, the prepared genome-wide siRNA library was primarily screened for siRNAs that suppressed NFAT5 nuclear translocation under high salt stress. During the screen, quality control was defined using the following criteria: in the reference control, the NFAT5 nuclear ratio in high salt condition was over 15% higher than that in isoosmotic condition and the nuclear ratio in ABL1 siRNA reference wells as the positive control was over 5% lower than that in control siRNA reference wells. The criteria were satisfied in all plates and siRNAs were ranked based on the value of NFAT5 nuclear ratio in each well. siRNAs that suppressed the ratio more than a positive control, ABL1 siRNA were defined as positive genes and 1291 candidate genes were obtained.

To implement a bioinformatics approach, KEGG enrich analysis of the positive genes were performed using an R package “ClusterProfiler” (ver.3.12.0).

### Immunofluorescence studies

After osmotic stress treatment, cells underwent the following fixation and staining steps: fixation by paraformaldehyde in PBS (3.7%), permeabilization and nuclear staining by 1% Triton X-100 and Hoechst 33258 in PBS, blocking by 5% skim milk, probing by the appropriate primary antibodies diluted by TBST, probing by the appropriate secondary fluorescent conjugated antibodies diluted by TBST, and replacing by PBS. Finalized assay plates were measured and analyzed using the CellInsight NXT with an optimized Cell Health Profiling BioApplication. In HES1 overexpression experiment, the transfected cells were seeded on cover glass coated with gelatin (Wako, Cat# 077-03155) in PBS and immunofluorescent staining were performed using the same protocol as mentioned above. After replacing by PBS, the cover grasses were mounted with Fluoromount (Diagnostic Biosystems, Cat#K024). Finalized samples were observed using a TCS SP5 confocal microscope (Leica Microsystems) and the captured images were analyzed using the CellInsight NXT with an optimized Cell Health Profiling BioApplication.

### Propidium iodide staining assay

Prior to capture images, the cells were incubated with culture medium containing Hoechst 33342 (Dojindo, Cat#346-07951, 1:1,000) and propidium iodide (PI; Dojindo, Cat#341-07881, 1:1,000) for 30 min in 5% CO2 at 37℃. After PBS substitution, the finalized plate was measured using the CellInsight NXT with an optimized Cell Health Profiling BioApplication.

### LDH release assay

LDH release was measured using the LDH-Cytotoxic Test (Wako, Cat#299-50601) basically following the manufacturer’s protocol. Briefly, culture medium was collected as medium sample. Cells were lysed using 0.1% Triton X-100 in PBS. Supernatants of centrifuged cell extracts were collected as lysate sample. Medium and lysate samples were individually mixed with reagents on microplates, and the absorbance was measured at 570 nm using a Varioskan Flash (Thermo Fisher Scientific) after around 5 min incubation at room temperature. Cell death ratio was calculated by LDH release (%) as follows: (absorbance (abs) of medium samples – background)/((abs of lysate samples – background) + (abs of medium samples – background)).

### Caspase 3 assay

Cells were lysed using 0.1% Triton X-100 in PBS. Supernatants of centrifuged cell extracts were mixed with caspase 3 substrate (Cayman CHEMICAL), PBS, DTT, and 2× Reaction Buffer (BioVision) in a black microplate and incubated at 37℃ for 2 h. Caspase activity was read at 400 nm/505 nm (Excitation/Emission) using the Varioskan Flash. The raw value divided by the protein concentration were normalized by each control sample under high salt condition as the Caspase 3 activity. Protein concentrations were measured using a protein quantification kit (Bio-Rad). Briefly, solutions A, B, and S were mixed according to the kit and combined with samples. Mixtures were then incubated for 30 min and absorbance was measured at 650 nm.

### Quantitative PCR (qPCR) analysis

Cells were treated with isoosmotic (≈ 300 mOsm/kg H2O) or high salt (≈ 400 mOsm/kg H2O) for 7.5 hours. Total RNA was isolated from cells using Isogen (Wako, Cat#319-90211) and reverse transcribed with ReverTra Ace qPCR RT Master Mix with gDNA Remover (Toyobo, Cat#FSQ-301), according to manufacturer’s instruction. Primers were designed using the Universal Probe Library Assay Design Center (Roche). Quantitative PCR was carried out using a LightCycler 96 (Roche) or QuantStudio 1 (Thermo Fisher Scientific) using FastStart Essential DNA Green Master (Roche, Cat#06924204001) or Kapa SYBR Fast qPCR Master Mix (Kapa Biosystems, Cat#KK4602). Data of each mRNA were normalized to that of RPS18 or GAPDH. Primer sequences are listed in Table S2.

### DNA microarray analysis

HeLa cells were reverse-transfected with 10 nM siRNA (control siRNA (Dharmacon, Cat#D-001210-04), NFAT5 siRNA (Dharmacon, Cat#D-009618-01), HES1 siRNA (Dharmacon, Cat#D-007771-08) for 48 hours. 7.5 hours after osmotic stress treatment (300 or 400 mOsm/kg H2O), total RNA was extracted from three independent wells of the same condition and mixed. RNA integrity was verified with a NanoDrop and an Agilent 2200 TapeStation. Biotinylated cDNA was prepared from total RNA using GeneChip WT PLUS Reagent Kit (Affymetrix) following the manufacturer’s instructions. Following fragmentation, single stranded cDNA was hybridized for 16 hr at 45°C on Clariom S Human Array Thermo Fisher Scientific, Cat#902927). Arrays were washed and stained in the GeneChip Fluidics Station 450 (Affymetrix) and were scanned using GeneChip Scanner 3000 7G. The data were analyzed with Affymetrix Expression Console Software 1.4.1 offered SST-RMA using Affymetrix default analysis settings. The data have been deposited in the NCBI Gene Expression Omnibus under accession number GSExxxxxx.

### ChIP-qPCR analysis

ChIP-qPCR analysis were performed using the SimpleChIP Chromatin IP Buffers and the SimpleChIP Enzymatic Cell Lysis Buffers A&B (Cell Signaling technology, Cat#14231, #14282) basically following the manufacturer’s protocol. Briefly, cells were fixed with formaldehyde, and chromatins were sheared with sonication into 150-900 bp DNA-protein complex. The complex was coprecipitated with NFAT5 antibody (SantaCrutz, Cat# sc-393887) or HES1 antibody (abcam, Cat#EPR-4226) and captured by Dynabeads Protein G (VERITAS, Cat#DB10004). The immunoprecipitated complexes were eluted and reverse cross-linked. The purified DNA fragment was analyzed using the same protocol of qPCR analysis as mentioned above with the primers listed in Table S2.

### Luciferase reporter assay

A fragment of human BGT1 promoter region (nucleotides −1732 to +22 of NM_003044.5) was cloned from a gDNA pool derived from HeLa cells and subcloned into pGL4.10[Luc2] (Promega, Cat#E665A). Two sites of NFAT5 binding motif (TGGAAANNYNY; Y is pyrimidine base) were manually detected and five sites of HES1 binding motifs were predicted using the JAPAR database (Fornes et al., 2019) in the region (Fig. 4G). The NFAT5 motif mutants (TaGAcANNYNY) and the HES1 motif mutant and deletion (Fig. 4J) were constructed from the wild-type BGT1 promoter and subcloned into pGL4.10[Luc2]. Dual-Luciferase reporter assay was performed using PicaGene Dual Sea Pansy Luminescence Kit (Wako, Cat# 301-05584) basically following the manufacturer’s protocol. Briefly, cells were co-transfected with the above BGT1 promotor constructs and pRL-TK (Promega, Cat# E224A). After osmotic stress treatment for 9 hours, cells were lysed in cell-lysis buffer in the kit. Cell extracts were centrifuged and supernatants were plated. PicaGene assay luminescence reagent and sea pansy luminescence reagent were dispensed on the plate and Firefly luciferase luminescence (FLU) from the pGL4.10 plasmids and Renilla luciferase luminescence (RLU) from the pRL-TK plasmids were sequentially measured using Varioskan Flash. Relative luciferase activity was calculated by dividing FLU by RLU.

### Quantification and statistical analysis

All data are represented as the mean ± SEM. Statistical tests were performed using R (ver. 4.2.1) with RStudio (ver. 2022.07.01). For all statistical analyses, *p < 0.05, **p < 0.01, and ***p < 0.001. Unpaired two-tailed Student’s t test or one-way ANOVA followed by Dunnett’s multiple comparisons test was used in this study.

## Supplementary Materials

Fig. S1. Validation of the genome-wide siRNA screen and the effects of HES1 knockdown on the nuclear translocation of endogenous NFAT5. Related to Fig. 1.

Fig. S2. A gamma-secretase inhibitor, DAPT, barely affected high salt stress-mediated HES1 induction, and exogenous HES1 overexpression did not affect GFP-NFATc3 nuclear translocation. Related to Fig. 2.

Fig. S3. The effect of HES1 knockdown on NFAT5-mediated gene induction and the cytoprotective interaction between HES1 and NFAT5 under high-salt conditions. Related to Fig. 3, 4 and 5.

Fig. S4. Validation of the BGT1 reporter vector and the requirement for DNA binding of HES1 to regulate NFAT5 activity. Related to Fig. 6

Data file S1. Positive Genes of Genome-wide siRNA Screen

Data file S2. Gene expression data of microarray analysis

## Acknowledgments

We thank all of the members of Laboratory of Cell Signaling in The University of Tokyo for critical discussions.

## Author contributions

Examples:

Conceptualization: H.R., I.N.

Methodology: H.R., Y.H., I.N.

Validation: H.R., Y.H.

Formal analysis:: H.R., Y.H.

Investigation: H.R., Y.H., I.N.

Visualization: H.R.

Funding acquisition: H.I., I.N.

Supervision: H.I., I.N.

Writing – original draft: H.R.

Writing – review & editing: H.I., I.N.

## Data and materials availability

All data are available in the main text or the supplementary materials.

