## Supplemental Figures for "HES1 contributes to high salt stress response as an enhancer of NFAT5-DNA binding"

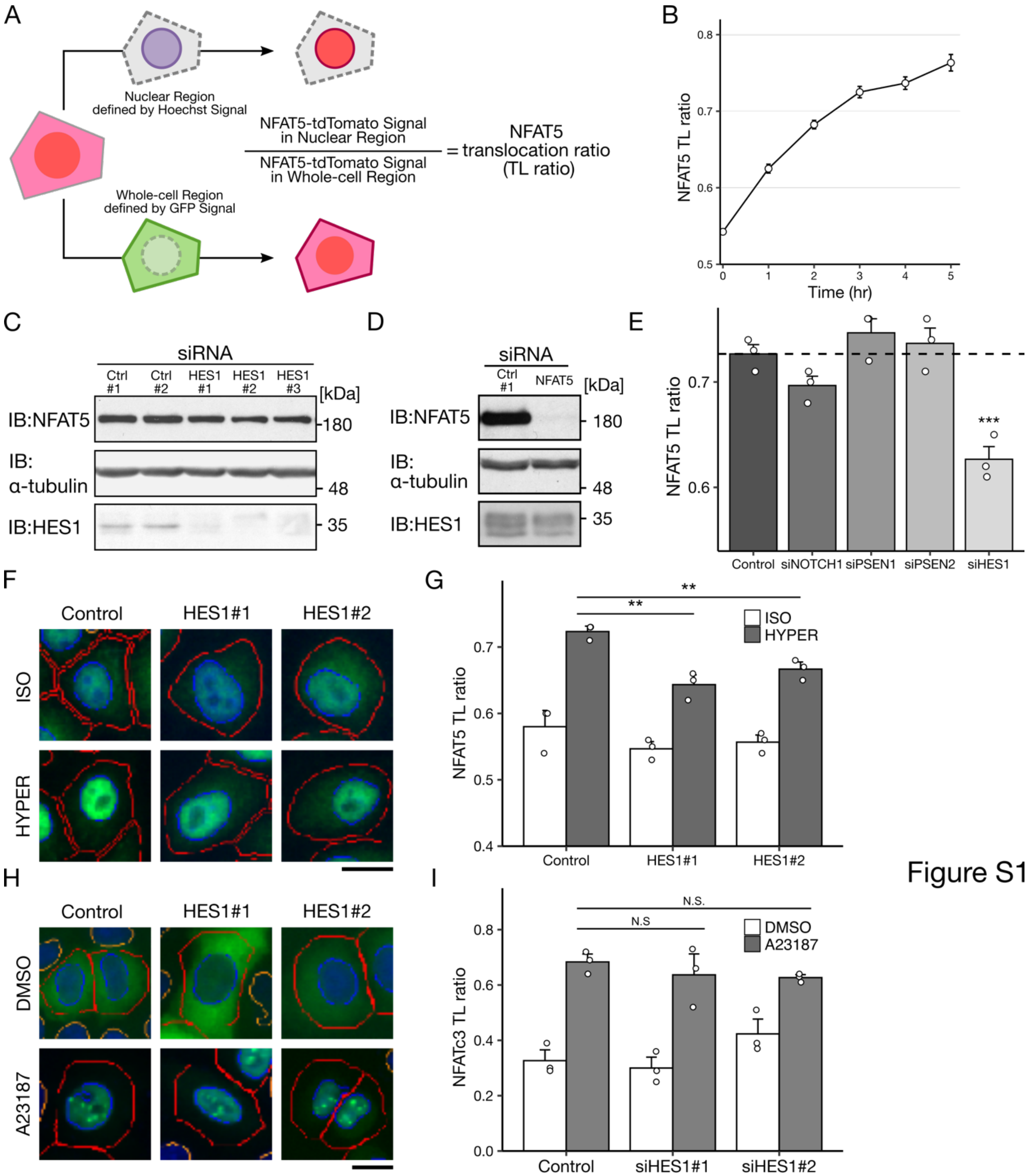


**Fig. S1. Validation of the genome-wide siRNA screen and the effects of HES1 knockdown on the nuclear translocation of endogenous NFAT5. Related to Fig. 1.** (**A**) A schematic model showing the design of the genome-wide siRNA screen to search for regulators of NFAT5 nuclear translocation. The fluorescence intensity of tdTomato fused with NFAT5Δ was measured in the nuclear and whole-cell regions defined by Hoechst and GFP signals, respectively. The NFAT5 translocation ratio (TL ratio) in each cell was calculated as follows: fluorescence intensity of tdTomato in the nuclear region/that in the whole-cell region. (**B**) Time course of the NFAT5 TL ratio calculated from the fluorescent images of NFAT5Δ-tdTomato HeLa cells (n = 3). (**C**) Efficacy of HES1 siRNAs on HES1 protein expression in HeLa cells. (**D**) Effect of NFAT5 siRNA on NFAT5 protein expression in HeLa cells. (**E**) NFAT5 TL ratio of the cells in which each Notch-related gene involved in the positive hit of the screen was depleted (n = 3). (**F**) Immunohistochemical analysis investigating the effect of HES1 depletion on endogenous NFAT5 nuclear translocation in HeLa cells. Green fluorescence showing endogenous NFAT5. Blue and red lines represent the border lines of the nuclear and whole-cell regions, respectively. (**G**) NFAT5 TL ratio calculated from the fluorescent images shown in (F) (n = 3, analyzing 700-1000 cells in each experiment). (**H, I**) Effect of HES1 depletion on GFP-NFATc3 nuclear translocation in HeLa cells. (H) Cells were treated with DMSO or a calcium ionophore, A23187, 20 nM, for 1 h. Blue and red lines represent the border lines of the nuclear and whole-cell regions, respectively. (I) NFATc3 TL ratio calculated from the fluorescent images shown in (H) (n = 3, analyzing 500-800 cells in each experiment).

In the bar graphs, individual values and the mean ±SEM are presented as white points and black bars, respectively. N.S., not significant; $^{**}p < 0.01$, $^{***}p < 0.001$. ISO, 300 mOsm; high Na, 500 mOsm (E) and 400 mOsm (F, G), 3 h. The scale bar represents 10 μm.

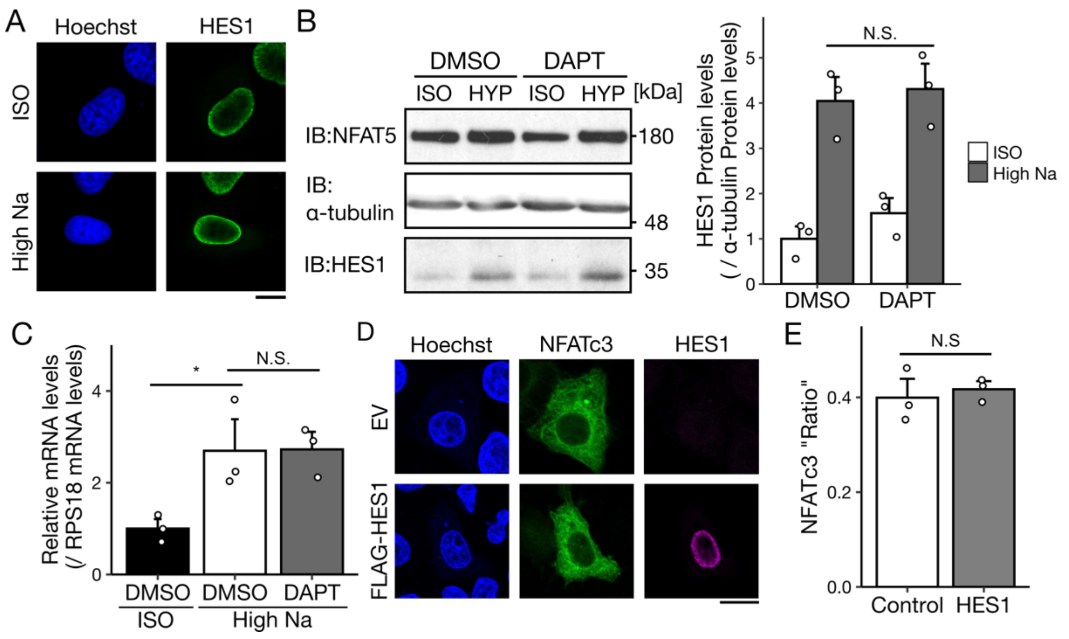


Figure S2

**Fig. S2. A gamma-secretase inhibitor, DAPT, barely affected high salt stress-mediated HES1 induction, and exogenous HES1 overexpression did not affect GFP-NFATc3 nuclear translocation. Related to Fig. 2.** (**A**) Immunohistochemical analysis of the subcellular localization of FLAG-HES1 in HeLa cells. (**B, C)** Effects of a gamma-secretase inhibitor, DAPT, on high salt stress-induced HES1 protein (B) and mRNA expression (C, n = 3). Cells were pretreated with DMSO or 10 µM DAPT for 30 min. (**D, E**) Effects of exogenous expression of FLAG-HES1 on GFP-NFATc3 nuclear translocation in HeLa cells (D). (E) NFATc3 TL ratio calculated from the fluorescent images shown in (D) (n = 3, analyzing 20-50 cells in each experiment). EV: empty vector, a negative transfection control.

In the bar graphs, individual values (white points) and the mean ± SEM are presented. N.S., not significant; $*p < 0.05$, ISO, 300 mOsm; high salt, 400 mOsm; 6 h in (B) and 5 h in (C). The scale bar represents 10 µm.

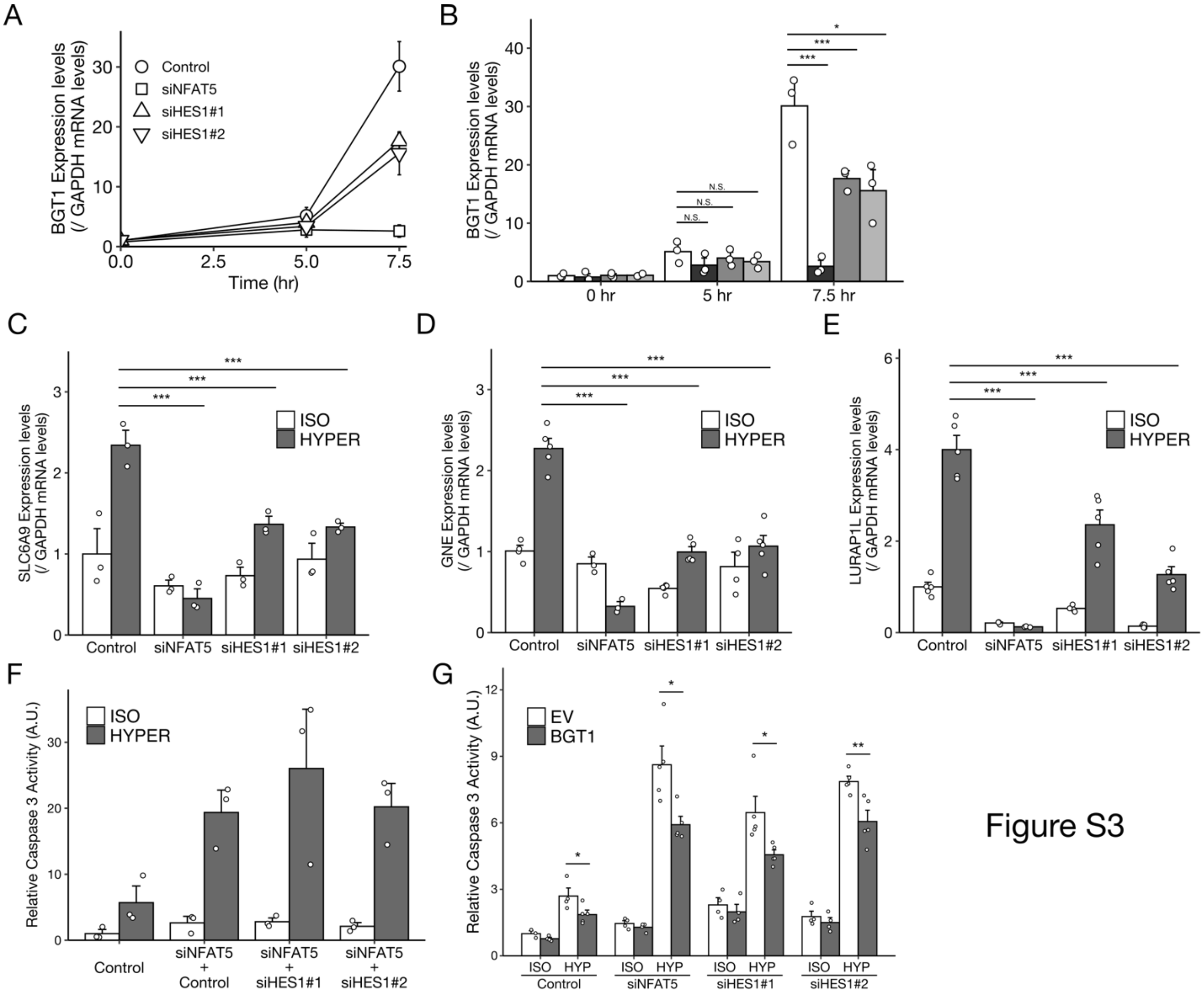


Figure S3

**Fig. S3. The effect of HES1 knockdown on NFAT5-mediated gene induction and the cytoprotective interaction between HES1 and NFAT5 under high-salt conditions. Related to Fig. 3, 4 and 5.** (**A**) Time course of BGT1 mRNA expression in HeLa cells quantified by qPCR (n = 3). (**B**) Effects of HES1 depletion on BGT1 induction under high salt conditions in HeLa cells (n = 3). The data sample is the same as in (A). (**C-E**) Effects of HES1 depletion on both NFAT5- and HES1-dependent genes detected in the microarray analysis, SLC6A9 (C, n = 3), GNE (D, n = 5) and LURAP1L (E, n = 5). (**F**) Effects of NFAT5/HES1 double knockdown on caspase 3 activity under hyperosmotic conditions (n=3). (**G**) An overexpression experiment on caspase 3 activity under high salt conditions, in which BGT1 was exogenously overexpressed in NFAT5- or HES1-depleted cells (n=5). EV: empty vector, a negative transfection control.

In the bar graphs, individual values (white points) and the mean ± SEM are presented. N.S., not significant; *p < 0.05, **p < 0.01, ***p < 0.001. ISO, 300 mOsm; high Na, 400 mOsm; (C-E) 7.5 h and (F, G) 9 h.

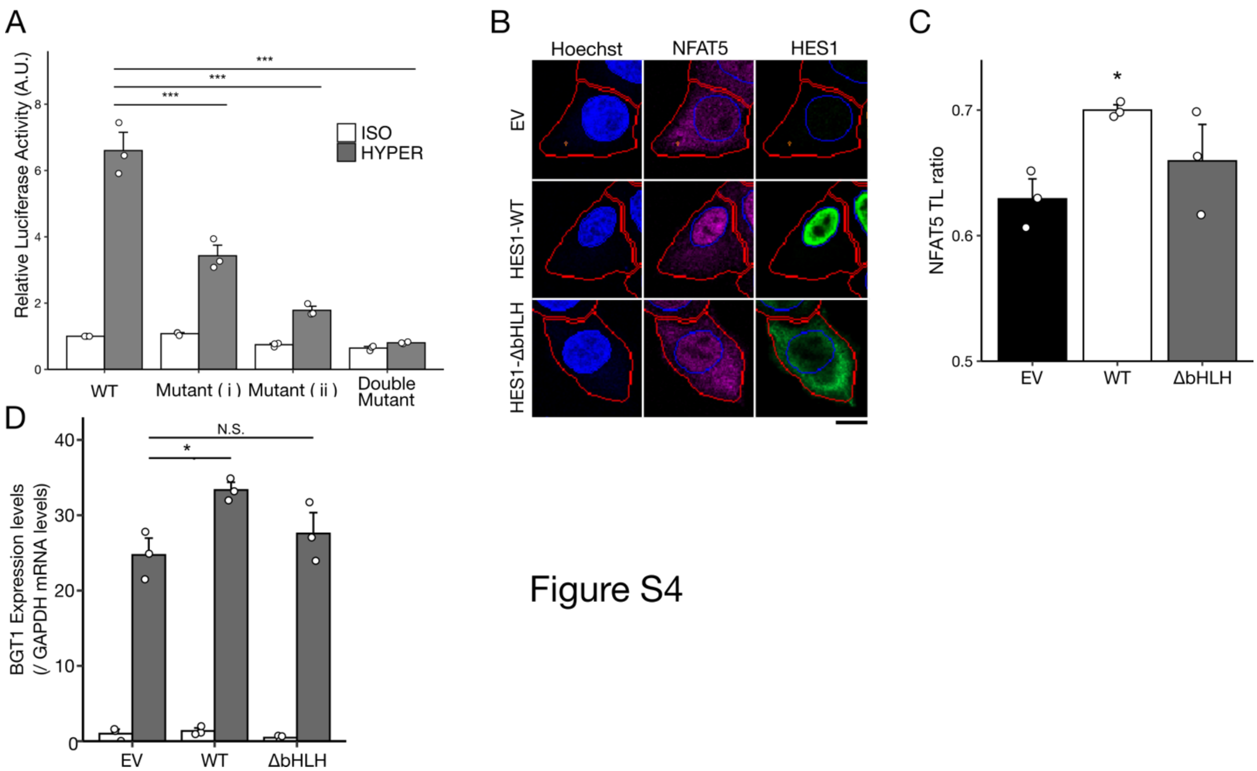


Figure S4

**Fig. S4. Validation of the BGT1 reporter vector and the requirement for DNA binding of HES1 to regulate NFAT5 activity. Related to Fig. 4.** (**A**) Contribution of NFAT5 binding motifs ((i) and (ii)) in the BGT1 promoter activity evaluated by point mutations (see Materials and Methods) ($n = 3$). (**B, C**) Effect of exogenous HES1 and a DNA binding-deficient mutant (HES1-ΔbHLH) on NFAT5 nuclear translocation in NFAT5Δ-tdTomato HeLa cells under isoosmotic conditions. The scale bar represents 10 µm (B). (C) NFAT5 TL ratio calculated from the fluorescent images shown in (B) ($n = 3$, analyzing 40-80 cells in each experiment). (**D**) Effects of exogenous HES1 and HES1-ΔbHLH expression on BGT1 mRNA expression under high salt conditions ($n=3$).
In the bar graphs, individual values (white points) and the mean ± SEM are presented. N.S., not significant; *$p < 0.05$, ***$p < 0.001$. ISO, 300 mOsm; high Na, 400 mOsm; 7.5 h in (A, D) and 3 h in (B, C).
